# Deep Learning Based Retrieval System for Gigapixel Histopathology Cases and Open Access Literature

**DOI:** 10.1101/408237

**Authors:** Sebastian Otálora, Roger Schaer, Oscar Jimenez-del-Toro, Manfredo Atzori, Henning Müller

**Author notes:** Both authors contributed equally to this manuscript (Sebastian Otálora, Roger Schaer).

## Abstract

Clinical practice is getting increasingly stressful for pathologists due to increasing complexity and time constraints. Histopathology is slowly shifting to digital pathology, thus creating opportunities to allow pathologists to improve reading quality or save time using Artificial Intelligence (AI)-based applications. We aim to enhance the practice of pathologists through a retrieval system that allows them to simplify their workflow, limit the need for second opinions, while also learning in the process. In this work, an innovative retrieval system for digital pathology is integrated within a Whole Slide Image (WSI) viewer, allowing to define regions of interest in images as queries for finding visually similar areas using deep representations. The back-end similarity computation algorithms are based on a multimodal approach, allowing to exploit both text information and content-based image features. Shallow and deep representations of the images were evaluated, the later showed a better overall retrieval performance in a set of 112 whole slide images from biopsies. The system was also tested by pathologists, highlighting its capabilities and suggesting possible ways to improve it and make it more usable in clinical practice. The retrieval system developed can enhance the practice of pathologists by enabling them to use their experience and knowledge to properly control artificial intelligence tools for navigating repositories of images for decision support purposes.

## 1. Introduction

Artificial Intelligence (AI)-based applications are quickly improving, and they are expected to be employed successfully in many fields, including medicine. In histopathology, AI tools may possibly modify the clinical practice in important global ways (for instance via computer aided diagnosis) or by supporting currently manual tasks (for instance via mitosis counting algorithms). Image retrieval on histopathology data is an intermediate solution that allows pathologists to enhance their clinical performance, while letting them the possibility to control the AI resources according to their experience and requirements. In recent years, various pathology departments have initiated a full digitization of their patient data and the stored samples[1]. Whole slide image (WSI) scanning now enables the on-screen visualization of high-resolution images from patient tissue slides. This opens the door to a wide range of image analysis and image navigation solutions [2], similar to what happened in the field of radiology, but with images of a much larger size in resolution up to 100, 000^2^ pixels. With fully digital workflows, pathologists obtain faster access to relevant patient data stored in such hospital repositories[2].

To fully exploit the potential of digital workflows, it is often necessary to structure the cases and all available metadata. Several pathology image platforms have already been proposed in the literature[3, 4, 5]. These platforms target either a specific research topic or have a flexible design to allow the inclusion of several data models and extensions for multiple applications [6]. The more recent platforms are web-based, which facilitates online sharing of large-scale imaging datasets and collaborative research[5]. Nevertheless, few of these platforms handle whole-slide imaging in an interactive and dynamic way, as the images are extremely large and not easy to visualize on-screen, some platforms are now moving towards large scale analysis of WSI. This limits the image analysis of WSIs, as it is routinely performed in the microscope by pathologists for the time being.

## 2. Methods

The retrieval system allows viewing whole slide images and searching for regions that are visually similar to manually defined regions of interest. Search is possible in a variety of data sources, including proprietary datasets that can be provided by the owner and public datasets (including for instance scientific literature). The workflow of the system is depicted in Figure1.

**Figure 1:**
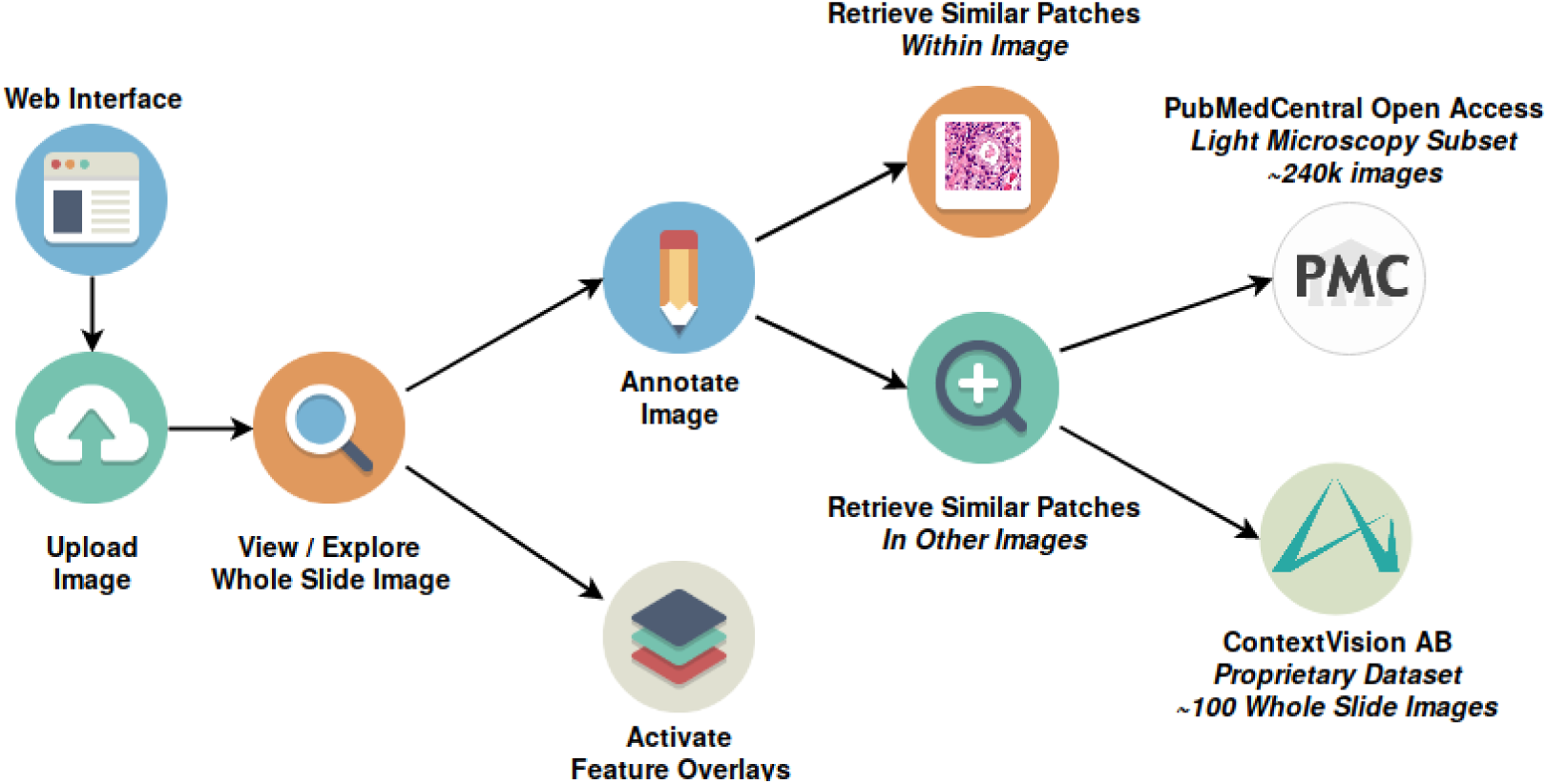
Workflow of interactions with the retrieval system

### 2.1 Programming Languages and Technologies

The viewer/annotation tool was developed as a “Node.js”^1^ application using the “Express”^2^ web framework, with JavaScript as the programming language for both front-and backend code, a set of python scripts were developed for deep feature extracion using a previously-trained model using the Keras framework^3^.

All data relative to annotations was stored using “CouchDB”^4^, a document-oriented database system designed for the web: it uses the standard JavaScript Object Notation (JSON) as a data format and the HyperText Transfer Protocol (HTTP) for communication. The retrieval system is based on a client-side only frontend developed with HyperText Markup Language 5 (HTML5) and JavaScript, communicating with a Java-based retrieval backend using REpresentational State Transfer (REST) web services.

Several existing libraries and software components were combined to develop the viewer/annotation tool, outlined below and in Figure2:¿.

### 2.2 Features

This section presents an overview of the main features that were implemented in the viewer/annotation tool, as well as in the retrieval system. It details the workflow of Figure1, of the user and illustrates how the system can help in fulfilling pathologists information needs.

**Figure 2:**
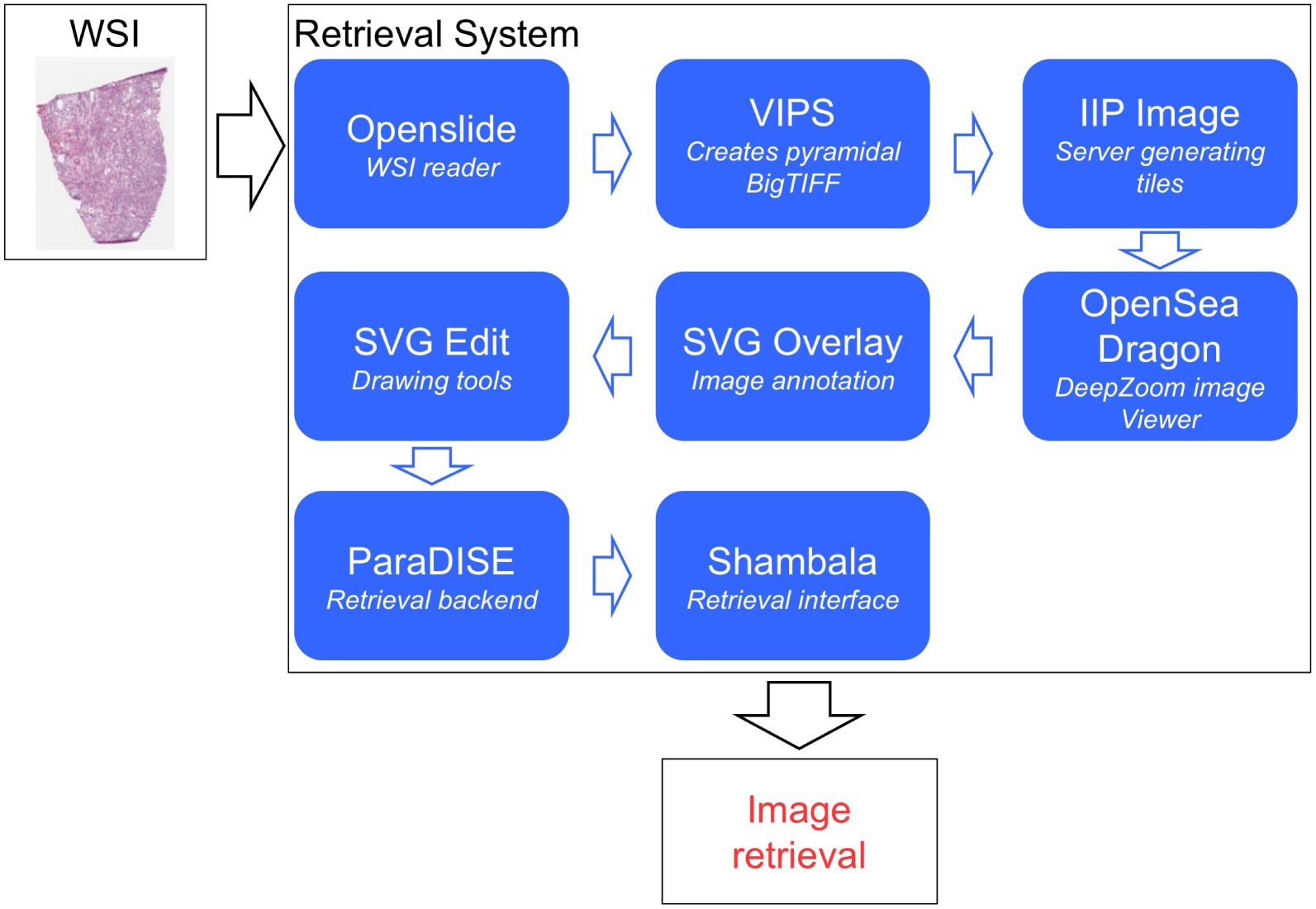
Libraries and software components used in the retrieval system

After the user has accessed the Web interface and logged in with his account, he can access existing datasets and images or upload a new image to the platform. Once the image processing process (described in the “Image Processing” section) has completed, he will be able to view the image and explore it by panning and zooming in on various regions of interest. From that point, the user has a choice to activate one or more “feature overlays”, that superimpose visual information on top of the original WSI, to highlight regions automatically segmented and classified by a deep learning algorithm, for example. To continue the workflow, a user would then typically create a new annotation within the image, by drawing an area of interest with one of several drawing tools (rectangle, ellipse, freehand, etc.). Once an annotation has been made, two possibilities are presented to the user:

- **Retrieve similar patches within the image:** When an annotation is selected, the system will automatically retrieve similar patches within the same image and present them to the user, who can then click on one of the results to navigate to that part of the image and inspect it more in detail
- **Retrieve similar patches in other images:** The user has the option to click on a link to open the retrieval interface that will allow him to perform a search of similar patches contained in other images. Two datasets are currently available to the user (see the “Datasets “section on page 20), which can be used for different types of retrieval tasks. Depending on the dataset, the user can then follow a link to a scientific article (e.g. PubMed Central) or navigate back to the viewer interface, visualizing a different image (e.g. images from a dataset of whole slide images available in the platform).

## 3. User Interface

This section presents an overview of the user interface that was developed and details each feature. The main user interface, shown in Figure 3, can be split into three principal areas: First, the navigation bar provides access to the various pages of the interface, including the homepage, a page with more information about the platform and a page allowing authorized users to upload new images to the platform. Secondly, a database/Feature selection & Annotation edition zone which allows switching between images contained in different datasets, as well as selecting feature overlays to display on top of an image (see Figure 4). In the same space also appears the annotation edition zone. Finally, in the viewer the main section where users can interact with images, zoom in and out quickly, create and edit annotations.

**Figure 3:**
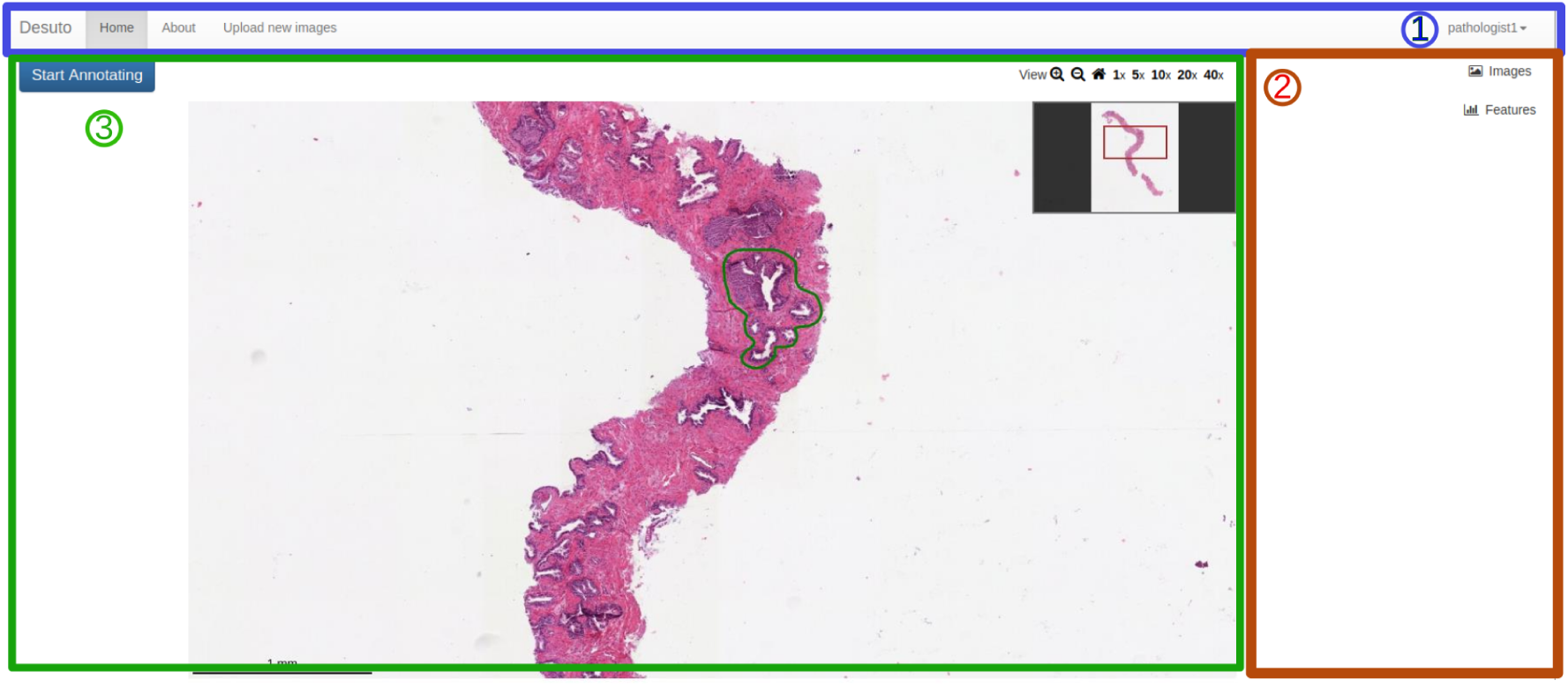
Overview of the main user interface. The three following sections are highlighted: (1) The navigation bar, (2) The main viewer container and (3) the database/feature selection panel and annotation edition zone.

**Figure 4:**
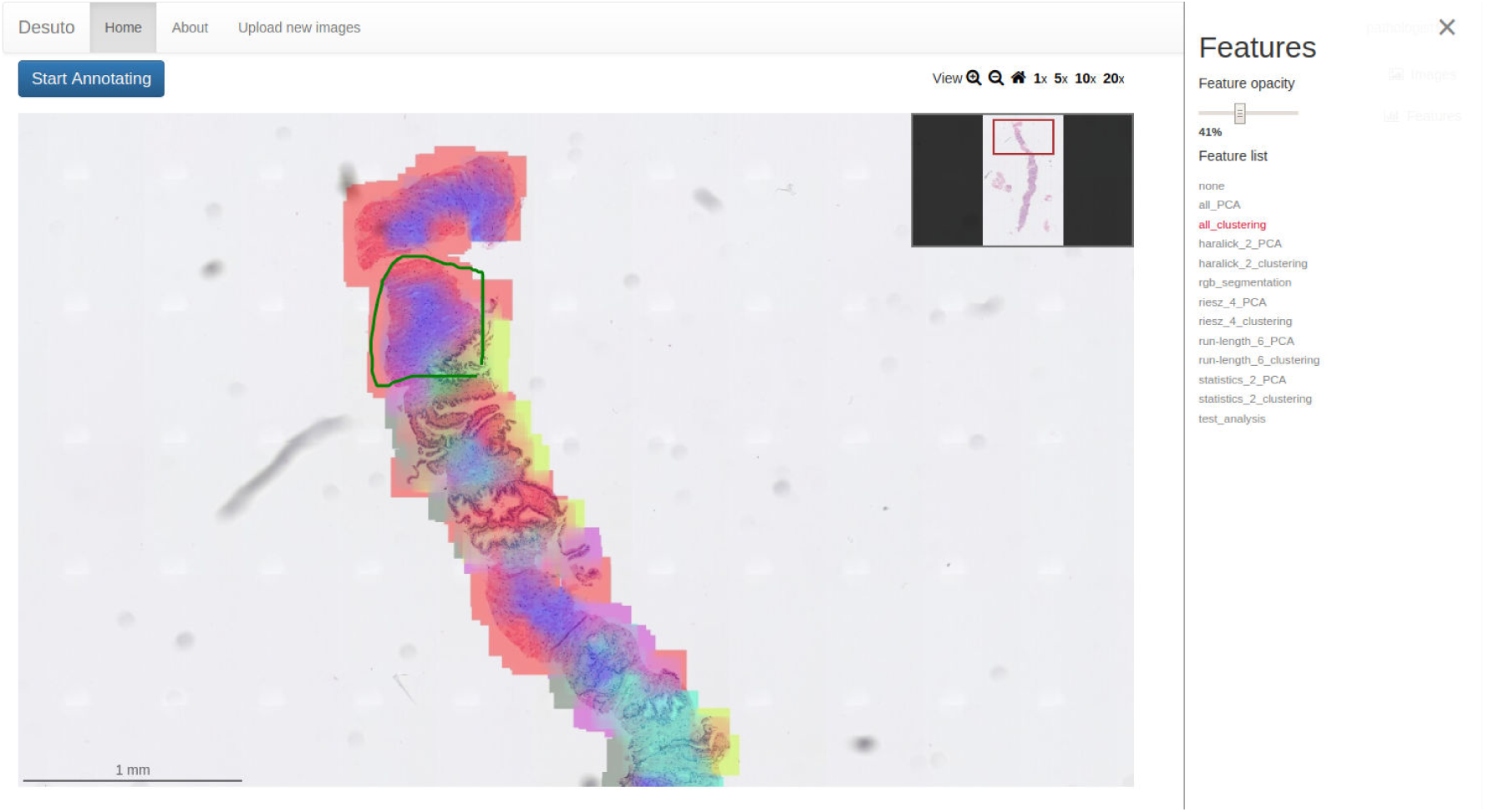
Coloured feature overlay of several types of computed features.

### 3.1 Retrieval Interface

The retrieval interface is tightly integrated with the viewer and opens in a pop-up window when the “Search for similar images” link is clicked in the annotation zone of a selected region (see Figure 5). It is composed of two main sections (see Figure6): The query section that contains a list of relevant query images and/or keywords at the left, and the results list which shows the retrieved results on the grid at the right. The retrieval interface is flexible and it supports multiple search modalities and options:

**Figure 5:**
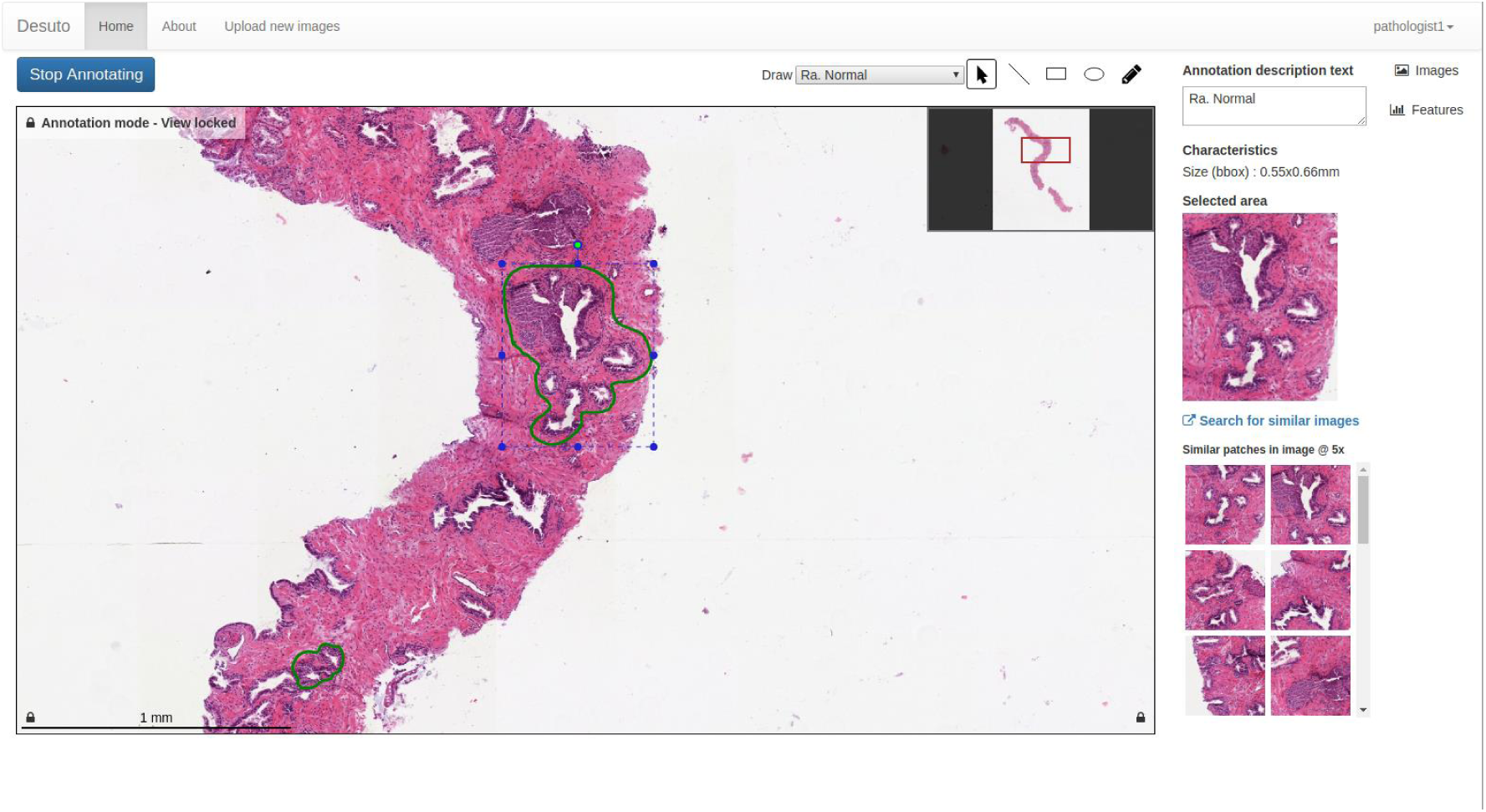
Preview of the selected annotation (bounding box) at the top and list of similar patches found within the same WSI at the bottom.

- **Content-based Image Retrieval (CBIR)**: By default, the region selected in the viewer appears in the “Relevant images” column in the retrieval interface, and visually similar patches are retrieved from the selected dataset. Any of the retrieved results can be dragged to the same column, in order to refine the search. By default, the visual similarity is computed using the histogram intersection of the feature vectors of the queries with the patches in the database, but other distance/similarity measures are supported as well.
- **Text-based retrieval:** If textual information regarding the query images or the search results (e.g. figure captions) is available, it is possible to add text search terms. This is the case in the example of the PubMedCentral dataset, where each image has an associated caption that can be used for finding more relevant results using keywords (see Figure 13).
- **Dataset and magnification options:** At the top of the results section are options enabling users to switch between the available datasets and different magnification levels within each dataset, if available (see Figure 14).

Advanced options such as the selection of individual visual features to use are also available, but where hidden for the user tests, to make the interface simpler and not overwhelm users with options. They can easily be shown or hidden depending on the targeted users (e.g. pathologists, computer scientists).

## 4. Evaluation

This section presents the datasets currently available in the system, as well as qualitative and quantitative evaluations of the system. They were performed via user test with pathologists and measuring the system’s retrieval performance based on ground-truth annotations. Evaluating a content-based histopathology retrieval system consist of two main components: The qualitative aspects of the system, such as interface usability, user experience, speed of the system; and the quantitatively aspects of the retrieval performance itself, i.e., how good is the system to retrieve relevant images from the system as measured in mean average precision and other performance measures.

### 4.1 Datasets

The main dataset used is a proprietary database of Contextvision AB (CVDB), including 112 WSIs of the prostate with manually annotated ROIs of healthy tissue and graded from thee to five according to the Gleason grading system[7, 8]. All the WSIs were loaded in the viewer and patches at distinct levels of magnification were extracted from the pyramidal content of the WSI (0.245 microns per pixel at the highest level) using the openslide library: 5X, 10X, 20X and 40X magnification patches were extracted from manually annotated ROIs. For each of the patches, both handcrafted Color and Edge Directivity Descriptor (CEDD) and deep learning features were extracted. This dataset highlights the capability of the retrieval system to manage proprietary datasets that may be provided by research groups or clinical pathology departments. The total number of patches used are showed in table1 and examples of the different magnifications are shown in Figure7.

**Figure 6:**
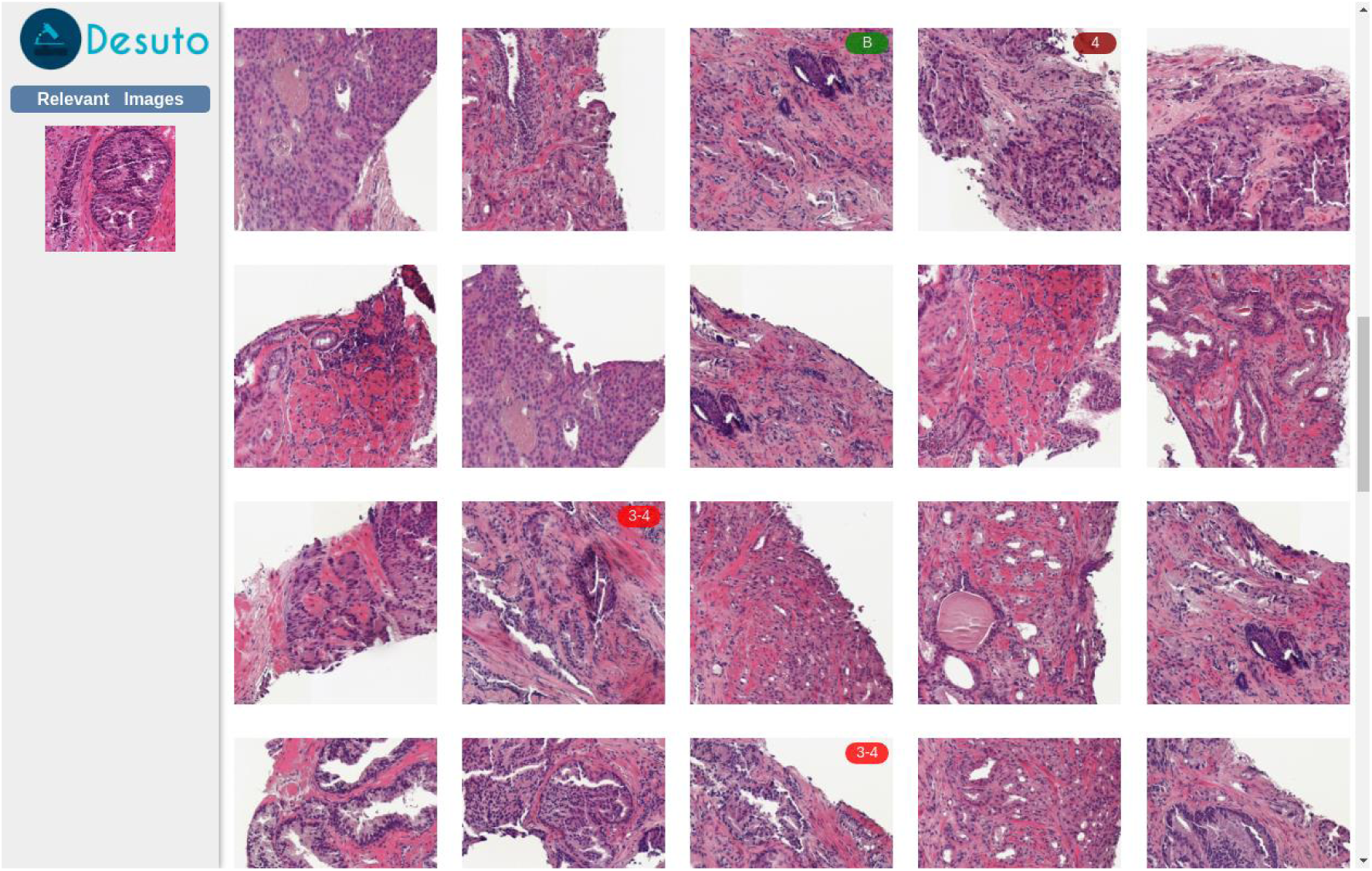
Preview of the selected annotation (bounding box) at the top and list of similar patches found within the same WSI at the bottom.

**Figure 7:**
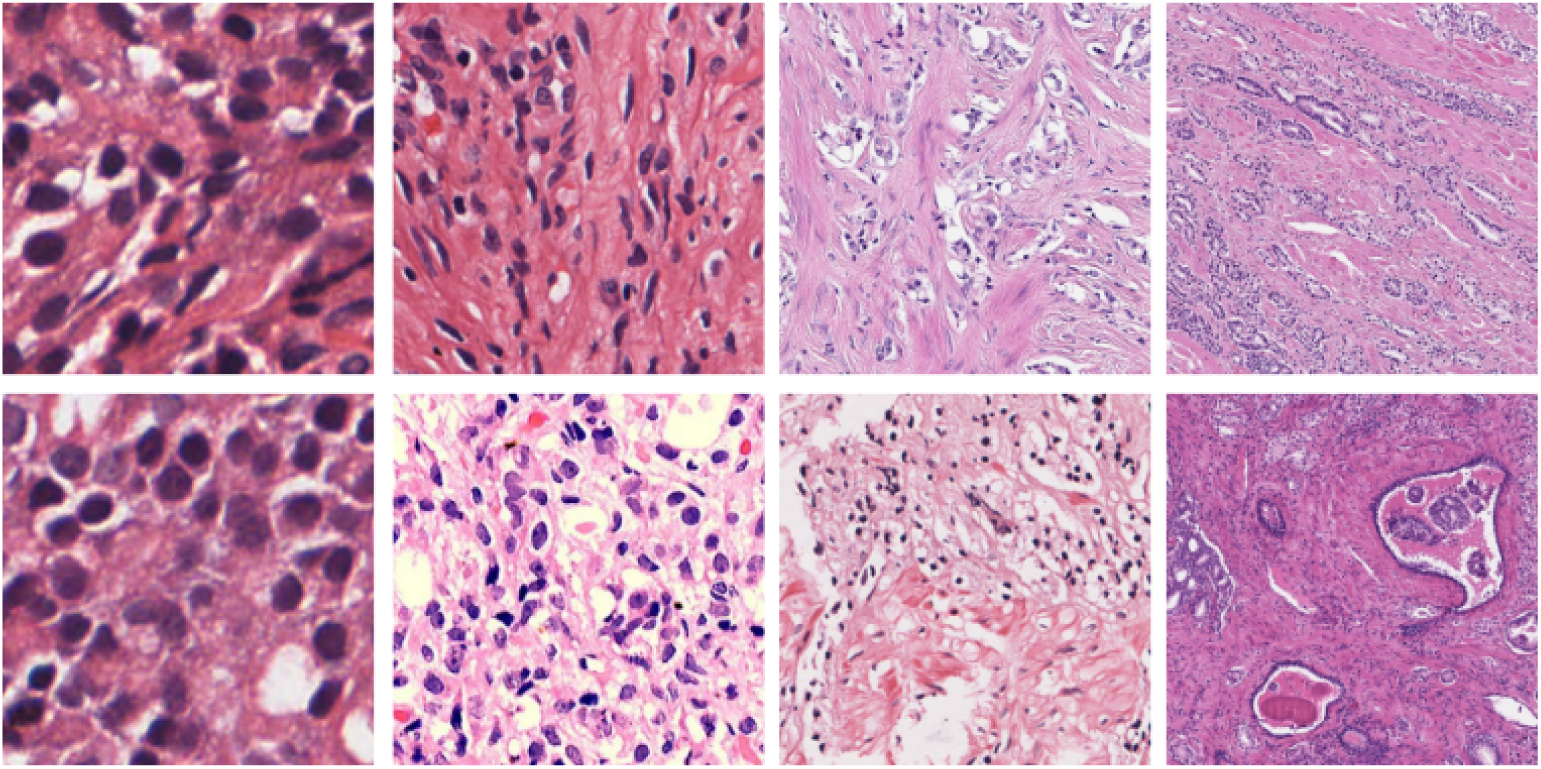
Example of patches from the CVDB dataset at WSI levels 0,1,2 and 3. The magnifications in these levels depends on the pixel resolution of the highest level which in the dataset is 0.2525 microns per pixel. The apparent magnifications corresponds to 40, 20,10 and 5X. Top row: patches from healthy ROIs; Bottom row: patches from ROIs with Gleason.

**Table 1:**
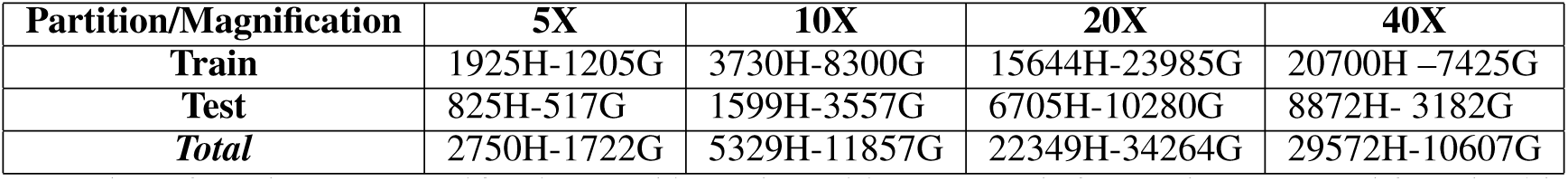
Number of patches per magnification used in each partition. H stands for patches extracted from healthy ROIs and G patches with some Gleason grades. In total our dataset consists of 118450 patches: 82913 for training/validation and 35537.

| Partition/Magnification | 5X | 10X | 20X | 40X |
| --- | --- | --- | --- | --- |
| **Train** | 1925H-1205G | 3730H-8300G | 15644H-23985G | 20700H-7425G |
| **Test** | 825H-517G | 1599H-3557G | 6705H-10280G | 8872H-3182G |
| **Total** | 2750H-1722G | 5329H-11857G | 22349H-34264G | 29572H-10607G |

The second currently available dataset consists of a subset of images from the PubMedCentral Open Access dataset which were classified as “light microscopy” images by an automatic classification algorithm[9]. This second dataset has very different characteristics, as it is not composed of WSIs (with patches extracted at various magnification levels), but rather of figures extracted from articles contained in a variety of medical journals. It has the added benefit of including a caption for each figure, enabling text-based search on this dataset, in contrast with annotation-level text only as in the case of WSIs. It includes around 240’000 images from the biomedical literature. This dataset highlights the capability of the retrieval system to manage publicly available collections of data, characterized by high variability.

### 4.2 Deep Learning Model and Features Indexing

For leveraging the discriminative power of deep learning models into our system, we trained a state–of–the–art architecture, the DenseNet model[10] which features a dense connectivity pattern among its layers. DenseNet introduces direct connections between any two subsequent layers with the same feature map size. The main advantage of DenseNet over other very deep architectures is that it reuses information at multiple levels without drastically increasing the number of parameters in the model, due to the inclusion of bottleneck and compression layers. The bottleneck layer applies a 1 × 1 convolution just before each 3 × 3 convolution to reduce the number of input feature maps. The compression layer uses a fraction of the feature maps in each transition layer.

**Figure 8:**
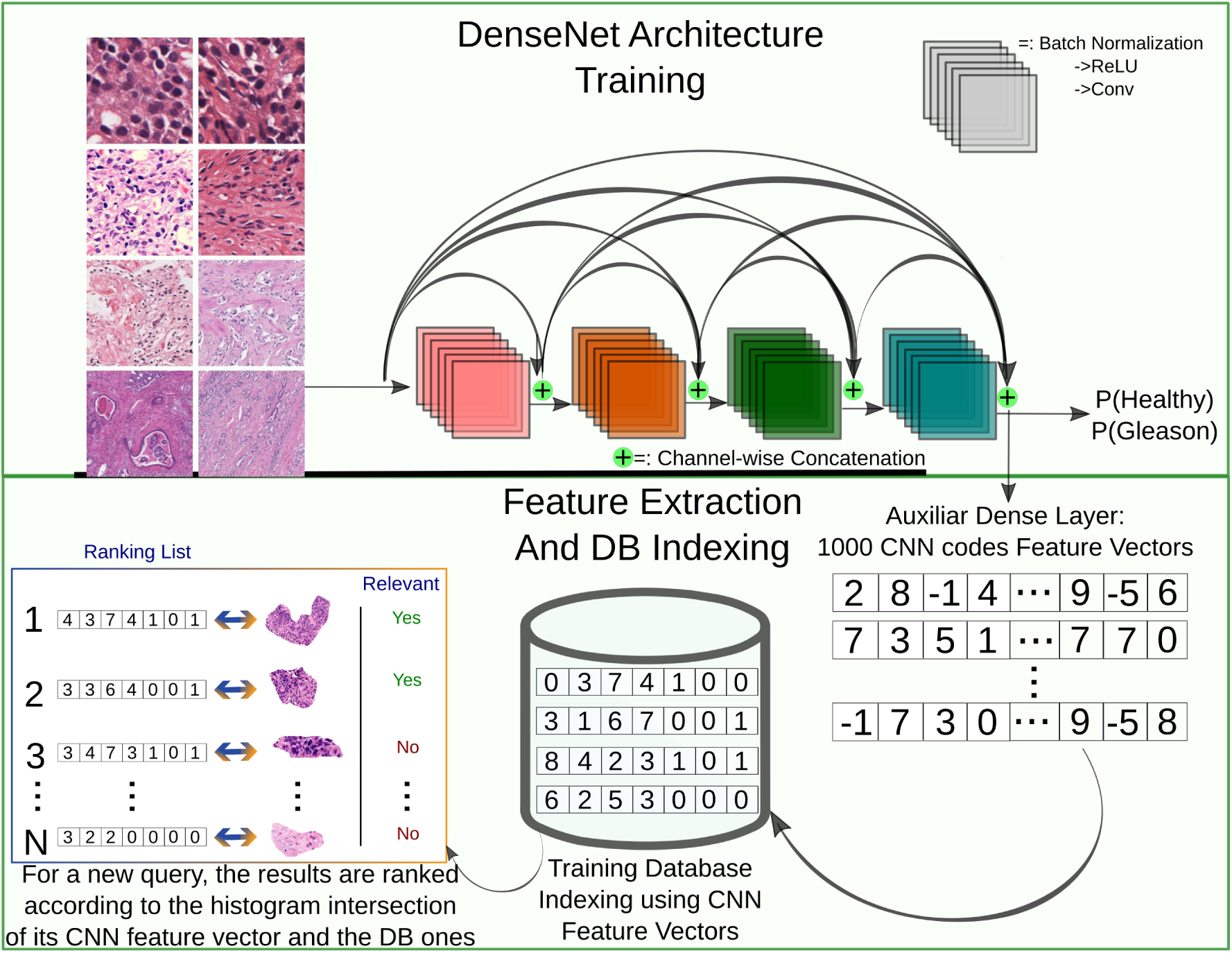
Schema of the DL model training and the subsequent feature extraction to build database (DB) index.

Once the network is trained, an auxiliary layer as shown in Figure8 is implemented, as the retrieval system computes similarities based on features vectors, it is thus necessary to extract feature vectors instead of having a probability of cancer/no cancer as output. The auxiliary layer is a dense layer which extracts 1000-dimensional feature vectors. After the feature vectors of all the patches are extracted, an index is built for each magnification level and then included in the ParaDISE retrieval engine.

### 4.3 Retrieval Performance

For assessing quantitatively the retrieval performance of our system, retrieval metrics from the US National Institute of Standards and Technology (NIST) evaluation procedures used in the Text Retrieval Conference (TREC^5^) were computed. The following five evaluation metrics were selected: mean averageprecision (MAP), geometric mean average precision (GM-MAP), binary preference (bpref), precision after ten patches retrieved (P10) and precision after 30 patches retrieved (P30). In Figure9 the precision–recall graph for both DL and CEDD features are compared. As has been shown in the literature, the deep-learning based representations outperform[11, 12] hand–crafted features such as CEDD in digital pathology tasks.

**Figure 9:**
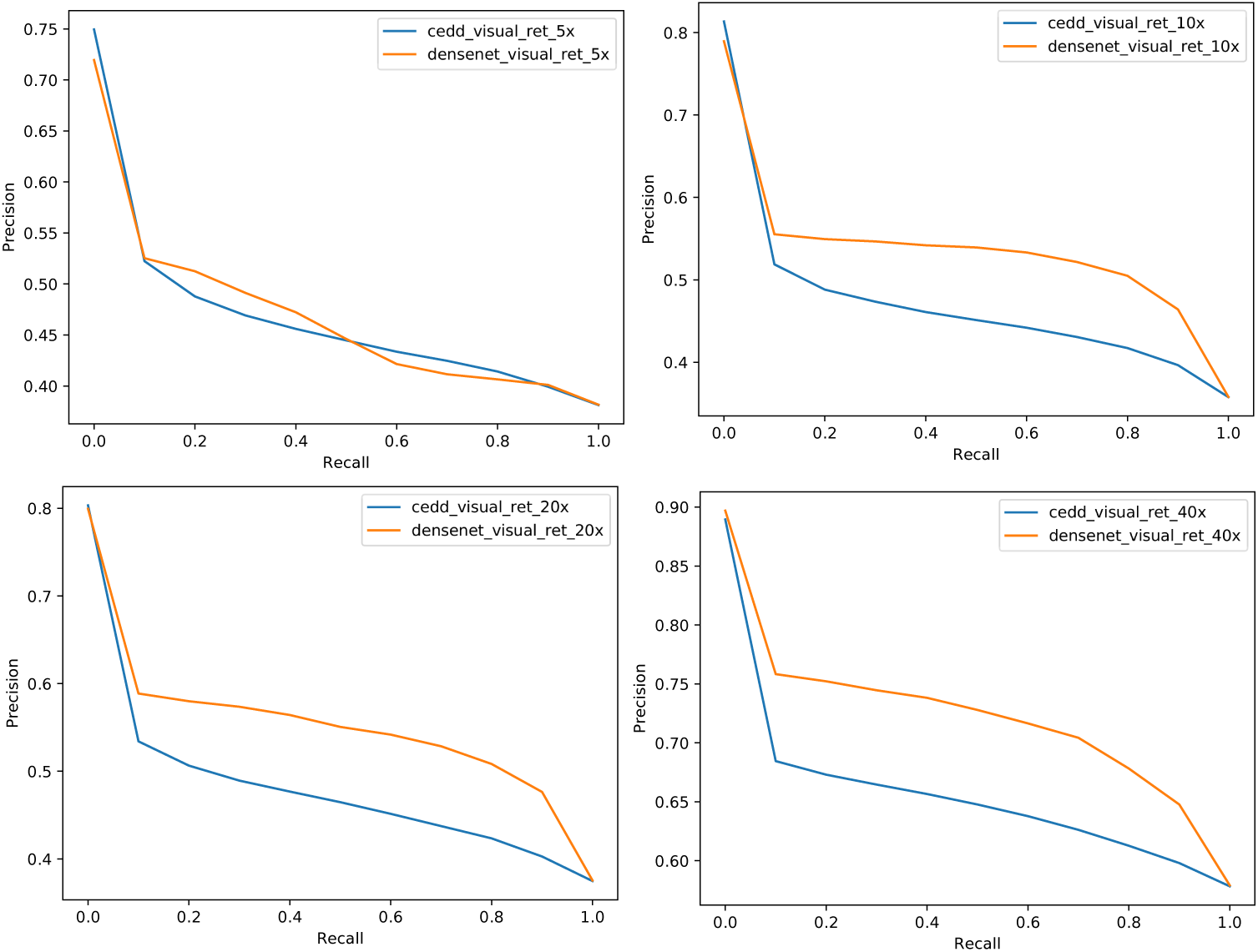
Precision-Recall graph of the retrieval performance of the visual features extracted using the DenseNet architecture (orange line) and the CEDD features(blue line) in four different magnification levels. 17’769 queries were performed over a subset of 8’292 training patches. The performance is similar for both features, nevertheless, the DL features are more stable and retrieve more relevant results.

### 4.4 User Tests

The interface of the system was evaluated qualitatively by two senior pathologists (P1, P2) who interacted with the system through a series of five tasks: (i) Marking a gland region and search for similar results in the WSI database, (ii) marking an area of inflammation, search in the PubMedCentral database, (iii) marking a Gleason 4 cribriform area and search for similar results in the WSI database, (iv) mark a Gleason 4 cribriform area and search for similar results in the PubMedCentral database and (v) free Interaction with the system. The pathologist remarked the ease of use of the different interfaces and the need for improving the quality of the results, since in some cases of PubMedCentral, the retrieved articles were not relevant for clinical use. The pathologist liked the organization of the different features and the fact that the more specific with the annotation they were, the more relevant images are found. Both of the pathologist remarked the need of more images to have a more robust results and model.

## 5. Conclusions

Clinical practice is getting increasingly tiring for pathologists due to growing complexity and time constraints. Interactive systems equipped with AI-based models could change the current work in histopathology. Differently from other applications of AI, such as computer aided diagnosis in radiology, digital pathology image retrieval is a solution that allows pathologists to enhance clinical performance by providing them a comprehensive set of tools for taking more informed decisions.

The developed retrieval system allows viewing whole slide images and searching for images similar to manually defined regions of interest in diverse data sources, including proprietary and public datasets. The use of proprietary datasets allows clinical departments and scientific research groups to develop their own resources to guide difficult diagnoses or to train students. The use of publicly available datasets, such as the PubMedCentral data allows pathologists to benefit from the increasing scientific knowledge available.

Searching the databases with both text search and content-based image retrieval opens new possibilities to pathologists. Searching by text and semantic concepts allows to search on the basis based on their own knowledge. Searching by visual content allows to search for similar cases, allowing the medical doctor to focus on visual aspects alone. The retrieval system can be helpful in complex situations, for instance when dealing with rare cases and it allows to save time in comparison to searching in books. Additionally, it represents an excellent way to train students. The system was tested by two pathologists, highlighting its capabilities and suggesting possible ways to improve it and make it better adapted to clinical practice.

The system was well accepted in the user tests. Comments were made regarding the retrieval quality and and regarding the limited number of indexed images: and for the described version of the system described here was already improved in terms of the retrieval results and amount of indexed data, was already improved integrating the comments of the both pathologists. Improving the data sources and further improving the performance of the system is currently the main target of our research group on this topic. In conclusion, the retrieval system presented in this paper can enhance the practice of pathologists. Nevertheless, merging the competences of pathologists with AI-based data analysis of large amounts of knowledge included in histopathology images is not an easy task and requires several cycles of user tests and feedback to be integrated into systems.

## Acknowledgement

This work was partially supported by the Eurostars project E! 9653 SLDESUTO-BOX and by Nvidia.

## Footnotes

1 http://nodejs.org as of February 19, 2018

2 http://expressjs.com as of February 19, 2018

3 https://keras.io/ as of February 19, 2018

4 http://couchdb.apache.org, as of February 19, 2018

5 https://trec.nist.gov/trec_eval/

## References

[1] Ronald S Weinstein, Anna R Graham, Lynne C Richter, Gail P Barker, Elizabeth A Krupinski, Ana Maria Lopez, Kristine A Erps, Achyut K Bhattacharyya, Yukako Yagi, and John R Gilbertson. Overview of telepathology, virtual microscopy, and whole slide imaging: prospects for the future. Human pathology, 40(8):1057–1069, 2009.

[2] Oscar Jimenez-del-Toro, Sebastian Otálora, Manfredo Atzori, and Henning Müller. Deep multimodal case–based retrieval for large histopathology datasets. In Patch–Based Techniques in Medical Imaging: Third International Workshop, Patch–MI 2017, Held in Conjuction with MICCAI 2017, Quebec City, Canada, September 14, 2017, Proceedings. Springer International Publishing, 2017.

[3] Raphaël Marée, Loïc Rollus, Benjamin Stévens, Renaud Hoyoux, Gilles Louppe, Rémy Vandaele, Jean-Michel Begon, Philipp Kainz, Pierre Geurts, and Louis Wehenkel. Collaborative analysis of multi-gigapixel imaging data using cytomine. Bioinformatics, 32(9):1395–1401, 2016.

[4] Claire McQuin, Allen Goodman, Vasiliy Chernyshev, Lee Kamentsky, Beth A Cimini, Kyle W Karhohs, Minh Doan, Liya Ding, Susanne M Rafelski, Derek Thirstrup, et al. Cellprofiler 3.0: Next-generation image processing for biology. PLoS biology, 16(7):e2005970, 2018.

[5] Daisuke Komura, Keisuke Fukuta, Ken Tominaga, Akihiro Kawabe, Hirotomo Koda, Ryohei Suzuki, Hiroki Konishi, Toshikazu Umezaki, Tatsuya Harada, and Shumpei Ishikawa. Luigi: Large-scale histopathological image retrieval system using deep texture representations. bioRxiv, page 345785, 2018.

[6] Kristian Kvilekval, Dmitry Fedorov, Boguslaw Obara, Ambuj Singh, and BS Manjunath. Bisque: a platform for bioimage analysis and management. Bioinformatics, 26(4):544–552, 2009.

[7] Brett Delahunt, Rose J Miller, John R Srigley, Andrew J Evans, and Hemamali Samaratunga. Gleason grading: past, present and future. Histopathology, 60(1):75–86, 2012.

[8] Jonathan I Epstein, William C Allsbrook Jr, Mahul B Amin, Lars L Egevad, ISUP Grading Committee, et al. The 2005 international society of urological pathology (isup) consensus conference on gleason grading of prostatic carcinoma. The American journal of surgical pathology, 29(9):1228–1242, 2005.

[9] Dimitrios Markonis, Roger Schaer, Alba García Seco de Herrera, and Henning Müller. The parallel distributed image search engine (paradise). arXiv preprint arXiv:1701.05596, 2017.

[10] Gao Huang, Zhuang Liu, Laurens Van Der Maaten, and Kilian Q Weinberger. Densely connected convolutional networks. In CVPR, volume 1, page 3, 2017.

[11] Geert Litjens, Clara I Sánchez, Nadya Timofeeva, Meyke Hermsen, Iris Nagtegaal, Iringo Kovacs, Christina Hulsbergen-Van De Kaa, Peter Bult, Bram Van Ginneken, and Jeroen Van Der Laak. Deep learning as a tool for increased accuracy and efficiency of histopathological diagnosis. Scientific reports, 6:26286, 2016.

[12] Angel Alfonso Cruz-Roa, John Edison Arevalo Ovalle, Anant Madabhushi, and Fabio Augusto González Osorio. A deep learning architecture for image representation, visual interpretability and automated basal-cell carcinoma cancer detection. In International Conference on Medical Image Computing and Computer-Assisted Intervention, pages 403–410. Springer, 2013.

